# Founder advantages in cell colony geometric organisation

**DOI:** 10.64898/2026.06.11.731426

**Authors:** Luke Honeybrook

## Abstract

Since the earliest microscopic observations, the geometric organisation of cells has captured biologists’ interest. Recent work by Gorgi et al. showed that bacterial colony organisation, including biofilms, can be explained across diverse species by radial expansion from fixed initial seeding sites and contact-inhibited growth, with little need for species-specific mechanisms. Here, we extend this geometric framework by incorporating seeding time as an additional driver of colony organisation. Using simulations and analytical models for expected colony size, we show that staggered seeding yields order of magnitude increases in the expected size of early seeded founder colonies. At realistic biofilm growth rates, a 2-day lag between founder and subsequent colony seeding produces an approximately 10-fold increase in expected founder size, while a 1-week lag produces a 25-fold increase. These findings provide a simple geometric basis for biological priority effects, illustrating temporal advantage alone can generate substantial spatial dominance, with implications for cardiovascular devices where host and bacterial cells compete in a ‘race for the surface’.

## Introduction

From the earliest microscopic observations, cellular life has been interpreted through geometry. Robert Hooke’s description of plant cells described their honeycomb-like architecture of pores and partitions, emphasising their spatial organisation as much as physiology [1]. This fascination with cell organisation has persisted across biology, from the competitive and cooperative organisations of single celled organisms to multicellular tissues and organs, whose origins and consequences remain central questions of biology. A longstanding question across these settings is what mechanisms produce the observed geometric organisations, and how generally they apply.

Recently, Gorgi et al. showed that the geometric organisation of diverse bacterial communities, surface biofilms at air-liquid interfaces, swimming and swarming populations, and in vivo host gut colonisation, is well described by a radial growth model [2]. In this model, colony boundaries grow radially outwards from initial seed locations, impinging at points of contact with other colony boundaries, and efficiently filling space to form Voronoi tessellations [3]. Complex species-specific mechanisms, such as reaction-diffusion kinetics [4], were found to be effectively unnecessary for predicting cell geometric organisations and that, surprisingly, predicting organisations required knowledge only of initial seed positions, colony growth rates, and little else. This framework has broad potential for improving understanding of how communities of cells form.

We extend this framework to quantify an important third factor: seeding times. In Gorgi et al.’s framework, all colonies were seeded, or began expanding, simultaneously [2]. In real biofilms, and across ecology more generally, community assembly is strongly dependent on priority effects, where species that establish early in an environment can come to dominate it [5]. Early biofilm colony compositions have been found to predict long-term population dynamics [6, 7] and to dominate infectious responses on hosts [8], which, within Gorgi et al.’s geometric paradigm, is consistent with the observation that founding colonies have far greater access to free space and can therefore outcompete colonies that arrive later [9]. However, the magnitude of this temporal effect, and its role in shaping final geometric organisation, remain to be quantified within the geometric framework that has proven successful for simultaneously seeded systems.

Priority effects are of direct clinical importance in medical device design, where competition for device surface space can determine patient outcome. Indeed, there is a ‘race for the surface’ between bacterial and host cells once devices are surgically implanted [10]. Bacterial biofilms establish on the surfaces of these devices, such as vascular grafts and ureteral stents, through a process of individual bacterial cells and small aggregates attaching to the surface, initiating microcolonies, expanding by clonal radial growth, and arrest when neighbouring colonies impinge [2, 9, 11-13]. Biofilms are typically far more challenging to clear than bacterial monolayers via antibiotics [11, 14]. Infections due to bacterial colonisation of cardiovascular devices are increasing in parallel with the rise of implantation, and frequently require surgical replacement or device explant that is associated with substantial morbidity and mortality [11, 14, 15].

A natural strategy for counteracting bacterial colonisation of cardiovascular devices is therefore to give the host’s own cells a temporal advantage in outcompeting bacteria for the available surface space—a priority effect. A vascular graft, for example, is considered ‘healed’ only if its luminal surface is fully covered by a host endothelial cell monolayer, yet complete coverage of current synthetic grafts is rare [15]. Accordingly, there has been much interest in encouraging endothelial tissue development in vivo through controlled host cell seeding, including the use of highly proliferative endothelial colony forming cells to outcompete bacterial colonisation [15, 16]. Strategies for controlling this endothelialisation and minimising biofilm formation may be better designed and understood through species-agnostic geometric principles of cell community organisation and the priority effects they entail.

Here, we extend Gorgi et al.’s geometric framework of bacterial community organisation [2] to incorporate the effect of seeding time. We used Monte Carlo simulations and analytical expressions for the expected size of individual colonies in both 2D and 3D and found that priority effects can manifest directly from spatiotemporal geometric considerations alone.

Indeed, two days of seeding time lag between the founder and subsequent colonies, at realistic biofilm growth rates, yielded an order of magnitude increase in expected founder colony size over the simultaneous seeding baseline. These results provide a quantitative basis for the founder advantage in biofilm formation and motivate seeding strategies that give host cells a temporal advantage needed to outcompete bacteria for surface space on cardiovascular devices.

## Materials and Methods

### Simulations of colony seeding and growth

Colonies were simulated as seeds undergoing radial, contact inhibited growth on a unit area (2D) or unit volume (3D) system with periodic boundary conditions. Each Monte Carlo instantiation had the same N prescribed seeding times, and for each seed a position was drawn uniformly at random over the entire domain at seeding time, in accordance with the Avrami model [17-21], which has previously been applied to a range of biological applications [22]. If a sampled position was within a covered region at the seeding time, the corresponding seed did not spawn, was recorded as a ‘phantom’ [19-21] and assigned a final size of zero in that run to ensure that the simulation samples spatially uncorrelated seeding. Periodic boundary conditions were implemented using wrapped distances, such that growth crossing one boundary re-entered from the opposite side.

Each instantiation therefore produces a vector of length N of colony size fractions evaluated at an analysis time τ. Repeating the run M times yields an M x N table of size fractions over independent instantiations. We use N = 100 colonies seeded from 0 hours to 120 hours mimicking similar in vitro experimental timescales [2]. The system size was set as 24 mm x 24 mm, corresponding to fields of view sizes used in similar in vitro experiments [2]. We prescribed a constant radial growth rate of 0.05 mm h^−1^ based on growth rates of *V. cholerae* biofilms [2]. We performed M = 10^4^ Monte Carlo instantiations. The MATLAB code used to perform these simulations is provided in the Supplementary Code files.

### Measures of colony size

We distinguish three distinct quantities, each of which may be referred to as a measure of colony size but which yield different results.

First is the typical colony area or volume (in 2D or 3D) distribution, which we generally refer to here as size, of a colony sampled uniformly from the tessellation [3]. It may be obtained by pooling all non-phantom entries from the entire M x N table of results into a single frequency histogram. In the infinite system and colony limit this yields a continuous probability distribution of colony sizes [23]. Equivalent results may also be obtained under alternative simulation conventions. For example, using a single very large instantiation in place of M independent instantiations, nonperiodic boundary conditions, or treating the total seed count itself as a Poisson random variable with mean equal to the prescribed expected number of seedings. These are valid if the system is large enough that boundary and finite colony size effects are negligible [3, 24, 25].

Second is the expected size of a colony labelled by its seeding time, obtained as the column average over all M rows, with phantoms included in the average. This is the unconditional expectation, E[X_i_], for colony i’s size labelled by its seeding time.

Third is the expected size of a colony conditional on it being non-phantom, obtained as the column average over all M rows but with zeros, instances where it was a phantom, excluded. This conditional expectation can be equivalently stated as E[X_i_ | X_i_ > 0]. These three quantities answer different biological questions and are conflated in some of the literature.

### Closed form expectations

The Avrami equation is a prediction of the expected total system fraction covered in the infinite system limit and is commonly written in a form with constant seed and growth rates J and G, respectively [17-21]. It is equivalently a sum over the infinite number of individual colony sizes, *E*[∑ *X*_*i*_]. In 2D for constant seed and growth rates, it has the form of Eqn. 1.

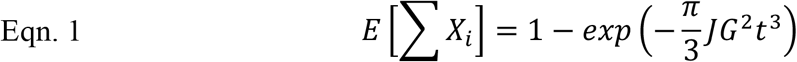

For the unconditional expectation E[X_i_] in finite systems with periodic boundaries we use the closed form expression derived in [26] for the expected size fraction of colony i at analysis time τ in any dimension, reproduced here as Eqn. 2.

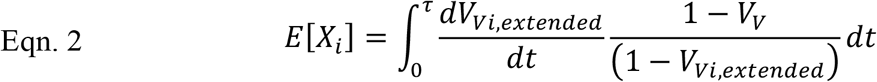

Where V_Vi,extended_ is the size colony i would achieve in the absence of other colonies, its ‘extended’ size [19-21]. V_V_ is the total covered fraction, a sum over all colonies, and may be calculated by 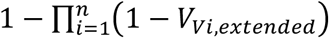: [24, 25]. In a finite periodic system, 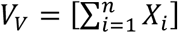 [24, 25].

From this unconditional expectation, the conditional expectation in finite systems with periodic boundaries may be derived via application of the conditional expectation formula, *E*[*X*_*i*_] = *E*[*X*_*i*_|*X*_*i*_ > 0]*P*(*X*_*i*_ > 0) [27]. Here, *P*(*X*_*i*_ > 0) is the probability that a chosen point in the system lies in uncovered space at the seeding time of colony i, which is 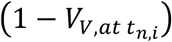:, yielding Eqn. 3.

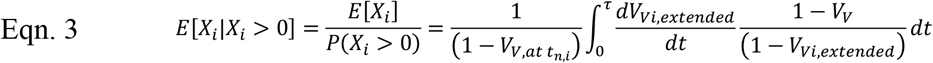

### Quantifying the founder advantage

The unconditional expectation of each colony’s size was evaluated numerically in 2D and 3D using Eqn. 2, with the same radial growth rate and system size as in the Monte Carlo simulations. The analysis time was set to the time of complete coverage (∼12 days). Two seeding regimes with N = 30 colonies were compared: 1) simultaneous seeding, with all colonies seeded at t = 0, 2) staggered seeding, with a single founder colony at t = 0 and the remaining colonies seeded at uniformly spaced times within a 5-day seeding window, offset from the founder by a prescribed lag time (Δt). The founder advantage was quantified as the fold-change of the founder’s expected size over the simultaneous seeding baseline.

## Results

### Measures of expected colony size are distinct

We first confirmed that the three measures of colony size defined above were distinct and that the closed form expressions reproduced the simulated distributions (Fig. 1B-D). For the typical colony size distribution, exact predictions in the infinite system limit are only available in special cases such as the 1D Avrami model with either constant seeding rate or simultaneous seeding [23]. More generally, approximate expressions have been derived. We plotted one such approximation, presented in [28], against the simulated typical colony size distribution in Fig. 1B and observed reasonable agreement.

**Figure. 1.**
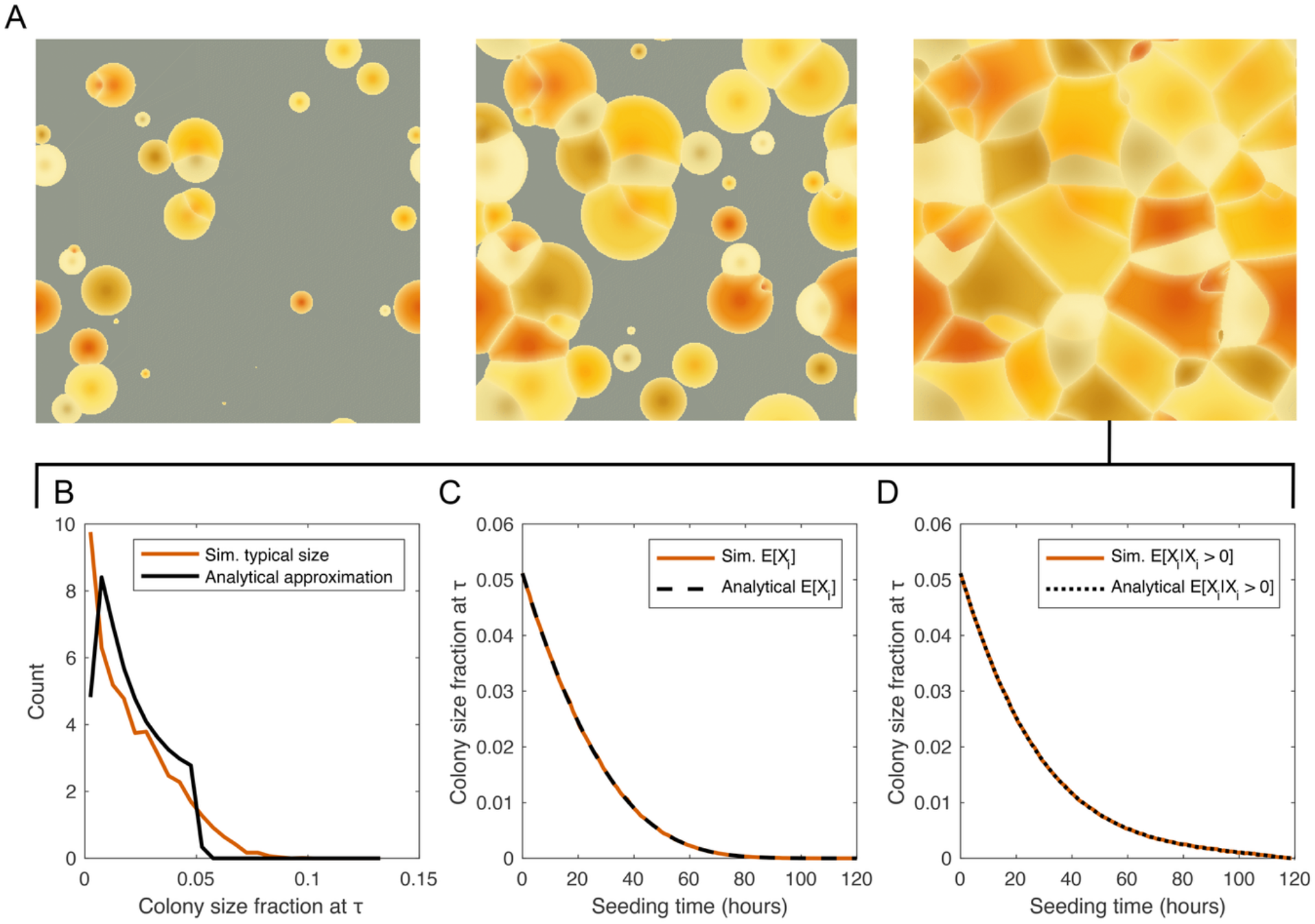
Simulations of staggered colony seeding and measures of expected colony size. A) Snapshots from a representative 2D simulation showing N = 100 colonies seeded at a constant seeding rate and undergoing radial contact inhibited growth. B) Typical colony size distribution: frequency histogram of pooled nonzero colony sizes evaluated at τ = 120 hours (red) compared with the prediction of an analytical approximation [28] (black). C) Expected colony size labelled by seeding time: averaged column means of colony sizes evaluated at τ = 120 hours (red) compared with the prediction of Eqn. 2 (black). D) Expected colony size labelled by seeding time excluding zeros: averaged column means of nonzero colony sizes evaluated at τ = 120 hours (red) compared with the prediction of Eqn. 3 (black).

For the unconditional expectation labelled by seeding time, Eqn. 2 accurately reproduced the simulated column mean across all seeding times (Fig. 1C, Table S1). For the conditional expectation, Eqn. 3 accurately reproduced the simulated column mean over nonzero results (Fig. 1D, Table S1). However, it departed strongly from the unconditional mean for later seeded colonies, inflating their expected sizes by ∼2-fold (Table S1). We also found that the unconditional expectation summed over all colonies recovered the Avrami covered fraction, whereas the sum over conditional expectations did not and could exceed unity (Fig. S1).

### The early founder advantage

We next used Eqn. 2 to quantify how the expected size of a colony depends on its seeding time. We found that when all colonies were seeded simultaneously, no colony had an advantage over any other across independent instantiations, and the expected size was identical for every colony in both 2D and 3D (Fig. 2A, C). The expected covered fractions of each colony were found to be equal to the inverse of the number of colonies seeded (1/N) in both 2D and 3D (Fig. 2A, C).

**Figure. 2.**
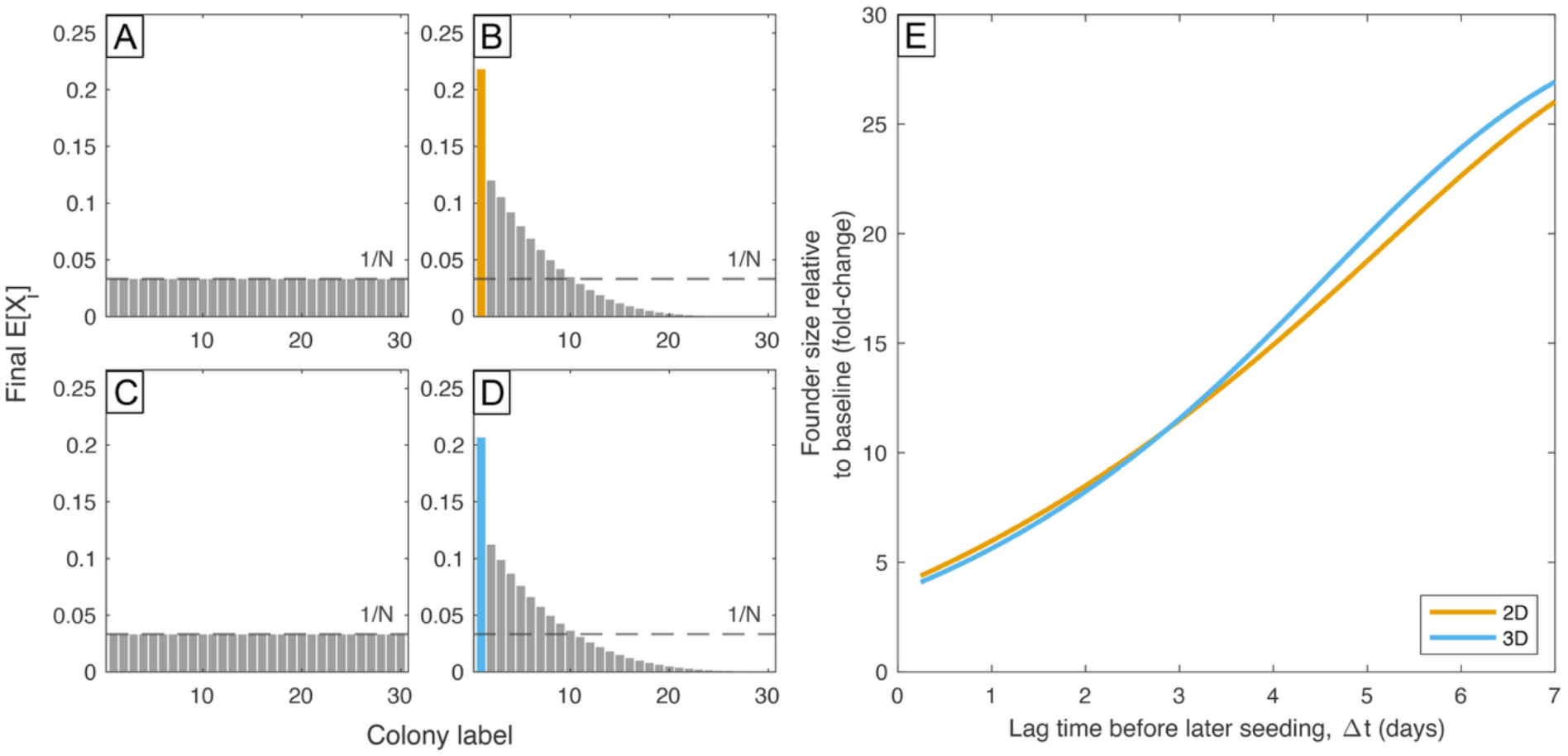
Founder advantage emerges when seeding times are staggered. A) Simultaneous seeding in 2D, all colonies achieve identical final expected size fractions of 1/N. B) Staggered seeding in 2D, the founder colony (orange) achieves a far greater final size compared to later seeded colonies (grey) and the simultaneous baseline of 1/N. C) Simultaneous seeding in 3D, all colonies achieve identical final expected size fractions of 1/N. D) Staggered seeding in 3D, the founder colony (blue) achieves a far greater final size compared to later seeded colonies (grey) and the simultaneous baseline of 1/N. E) Expected size of the founding colony, relative to simultaneous seeding baseline, as a function of lag time between the founder and the next earliest seeded colony in 2D (orange) and 3D (blue).

Staggering the seeding times produced a markedly different result. With a simple time lag between the first seeded colony (the founder colony) and later seeded colonies of 1 day, the founder acquired an expected final size 5-fold larger than the simultaneous seeding baseline (Fig. 2B, D). With a lag time of 2 days between the founder and subsequent seedings, the founder was approximately 10-fold larger, an order of magnitude increase, than the simultaneous seeding baseline. At a lag time of 1 week, the founder colony was approximately 25-fold larger than the simultaneous seeding baseline (Fig. 2E). The founder advantage scaled approximately linearly with lag time and was very similar in both 2D and 3D scenarios (Fig. 2E).

## Discussion

We have distinguished three distinct measures of colony size. The first is the typical colony size distribution, the most extensively studied measure of size in classical work on Voronoi tessellations [3]. It describes the distribution of sizes one would observe across all instantiations and, because seed locations are uniformly random within each instantiation, yields a heterogeneous distribution in which some colonies achieve much larger sizes than others [23]. Crucially, this heterogeneity is intrinsic to the random spatial layout of the seeds and arises regardless of seed time synchrony. Indeed, both simultaneous seeding and staggered seeding produce distributions with strong heterogeneity [3, 23]. This measure is therefore strongly confounded by spatial positioning effects and cannot easily isolate spatiotemporal priority effects.

The unconditional expectation of a colony’s size labelled by its seeding time, E[X_i_], averages out the confounding effect of spatial randomness across independent samples and so isolates the pure effect of seeding time on expected colony size. Closed form expressions for E[X_i_] in any dimension were recently derived [26] and we show here that the sum of this measure over all individual colony sizes accurately recovers the Avrami prediction. This is due to the linearity of expectation identity, 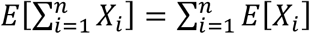 [27].

The third measure we investigate is the conditional expectation of seed time-labelled colony size given that the colony was successfully seeded in a given instantiation, E[X_i_ | X_i_ > 0], which we derive here for the first time. An expression of the same functional form as Eqn. 3 was presented in [29] but was suggested to be a measure of typical colony size that would recover the Avrami prediction upon summation. We show that a function of this form is in fact a conditional expectation and does not recover the Avrami prediction when seeding times are staggered. Indeed, its sum over all colonies can exceed unity. To better understand why the conditional expectation can be useful but is not considered further here, requires a discussion of the different methods of random seeding.

We considered here the case of uniform random seeding, consistent with previous models of biofilm formation that specify random seeding [2, 4, 9]. When colonies are seeded simultaneously this produces a Voronoi tessellation, but when seeding times are staggered the result is termed an additively weighted Voronoi tessellation [3]. The traditional model for producing additively weighted Voronoi tessellations is via uniform random seeding across the entire system, including occupied regions, and treating seeds that fall within occupied space as phantoms—spatially uncorrelated seeding [3, 17-21]. An alternative random seeding strategy is possible, whereby seeds are chosen from uniform random positions in unoccupied regions only—an example of spatially correlated seeding. The use of spatially uncorrelated seeding for the generation of additively weighted Voronoi tessellations has become convention partly because it is simpler to model analytically but also because it guarantees that, on average, a prescribed constant seeding rate per unit of unoccupied system is maintained over time [30]. It is worth noting that the distribution of E[X_i_], despite being a quantity derived for spatially uncorrelated seeding, yields a good approximation of the typical colony size distribution in a spatially correlated seeding scenario [26].

In the context of colony formation and their geometric organisation, the unconditional and conditional expectations may both be reasonably considered as useful in addressing questions of priority effects. However, because conventional models of Voronoi tessellations that are applied to colony formation [2, 4], and across biology more generally [22], assume spatially uncorrelated seeding [3], we suggest the unconditional expectation is the more appropriate measure.

We found that under simultaneous seeding at uniform random positions the unconditional expected sizes of all colonies across independent samples are equal, in stark contrast to the typical colony size distribution, which is strongly heterogeneous [3]. We further found that the expected sizes upon complete system coverage equalled the inverse of the number of colonies seeded in both 2D and 3D. The arithmetic mean of the typical colony size distribution in the infinite limit recovers the same value—the inverse of the expected number of colonies per unit region [3, 23]. This occurs because both quantities have the same sum and number of set elements and therefore must have the same arithmetic mean.

We next found that when colony seeding times were staggered, and in particular when a time lag was introduced between the founder colony and subsequent colonies, the expected size of the founder colony increased dramatically over the simultaneous seeding baseline, by up to an order of magnitude. Indeed, when subsequent colonies were delayed in arrival by 1 week, the founder colony achieved a ∼25-fold increase in expected size. This provides a physically meaningful underpinning of priority effects, observed across microbial ecology [5, 8] and during direct observations of biofilms [6, 7, 9], in the context of spatiotemporal competition [2].

The fold-change founder advantage was similar in 2D and 3D, scaling approximately linearly with lag time in both settings. This dimensional insensitivity may be useful for translational interpretation. In vitro biofilm studies are commonly performed on flat 2D culture surfaces [6, 7, 13], whereas medical device surfaces, including vascular grafts, are increasingly being developed from porous synthetic and tissue engineered materials that permit 3D proliferation [15]. The finding that the expected founder size advantage is comparable across 2D and 3D systems suggests that conclusions from conventional 2D culture models in the context of spatiotemporal priority effects may be more readily transferable to 3D settings. This contrasts with many other cell culture models where dimensionality can substantially alter cell behaviour and weaken relevance to in vivo settings [13].

## Conclusion

The geometric organisation of colony-forming cells, including bacterial biofilms, is well described by Voronoi tessellations arising from radial, contact inhibited clonal expansion from initial seeding sites. We extend this framework to incorporate seeding time and quantify the founder advantage of early seeded colonies. Staggered seeding produced a strong founder advantage in expected colony size, similar in 2D and 3D, and scaling approximately linearly with lag time. Just two days between the founder and subsequent colonies, at realistic biofilm growth rates, yielded an order of magnitude increase in expected size. These results show that the priority effects observed across microbial ecology and biofilm dynamics can manifest directly from simple spatiotemporal geometric considerations. They further suggest that surface seeding strategies could exploit even modest temporal advantages to favour host coverage over bacterial colonisation in the ‘race for the surface’ on cardiovascular devices.

## Acknowledgements

British Heart Foundation grant FS/4yPhD/F/20/34134 (LH).

## Data Availability Statement

MATLAB code used to produce Figure 1, S1, and Table S1 are provided in the Supplementary Code files.

## References

1. Hooke, R., Micrographia. 1665, London, England: The Royal Society.

2. Gorgi, M., et al., Geometric ordering in bacterial communities. Proceedings of the National Academy of Sciences, 2026. 123(20): p. e2526643123.

3. Okabe, A., et al., Spatial Tessellations: Concepts and Applications of Voronoi Diagrams. 2nd ed. 2000, Chichester, England: John Wiley & Sons.

4. Chacón, J.M., W. Möbius, and W.R. Harcombe, The spatial and metabolic basis of colony size variation. The ISME Journal, 2018. 12(3): p. 669–680.

5. Fukami, T., i>Historical Contingency in Community Assembly: Integrating Niches, Species Pools, and Priority Effects. Annual Review of Ecology, Evolution, and Systematics, 2015. 46(Volume 46, 2015): p. 1–23.

6. von Bronk, B., A. Götz, and M. Opitz, Locality of interactions in three-strain bacterial competition in E. coli. Physical Biology, 2019. 16(1): p. 016002.

7. Eigentler, L., F.A. Davidson, and N.R. Stanley-Wall, Mechanisms driving spatial distribution of residents in colony biofilms: an interdisciplinary perspective. Open Biology, 2022. 12(12): p. 220194.

8. Devevey, G., et al., First arrived takes all: inhibitory priority effects dominate competition between co-infecting Borrelia burgdorferi strains. BMC Microbiology, 2015. 15(1): p. 61.

9. Eigentler, L., et al., Founder cell configuration drives competitive outcome within colony biofilms. The ISME Journal, 2022. 16(6): p. 1512–1522.

10. Gristina, A.G., Biomaterial-Centered Infection: Microbial Adhesion Versus Tissue Integration. Science, 1987. 237(4822): p. 1588–1595.

11. Francolini, I., L. Hall-Stoodley, and P. Stoodley, Biofilms, Biomaterials, and Device-Related Infections, in Biomaterials Science, W.R. Wagner, et al., Editors. 2020, Academic Press. p. 823–840.

12. Matoz-Fernandez, D., et al., Comment on “Rivalry in Bacillus subtilis colonies: enemy or family?”. Soft Matter, 2020. 16(13): p. 3344–3346.

13. Hunter, L., et al., Cell Calcification Models and Their Implications for Medicine and Biomaterial Research. Advanced Healthcare Materials, 2026. 15(6): p. e01104.

14. Elek, S.D., Experimental Staphylococcal Infections in the Skin of Man. Annals of the New York Academy of Sciences, 1956. 65(3): p. 85–90.

15. Seidman, M.A., R.F. Padera, and F.J. Schoen, Cardiovascular Medical Devices: Stents, Grafts, Stent-Grafts and Other Endovascular Devices, in Biomaterials Science, W.R. Wagner, et al., Editors. 2020, Academic Press. p. 1033–1050.

16. Melero-Martin, J.M., Human Endothelial Colony-Forming Cells. Cold Spring Harbor Perspectives in Medicine, 2022. 12(12): p. a041154.

17. Kolmogorov, A.N., On the Statistical Theory of Crystallization of Metals [in Russian]. Izv. Akad. Nauk SSSR, 1937. Ser. Mat.(3): p. 355–359.

18. Johnson, W. and R. Mehl, Reaction kinetics in processes of nucleation and growth. Trans. Metall. Soc. AIME, 1939. 135: p. 416–442.

19. Avrami, M., Kinetics of Phase Change. I General Theory. The Journal of Chemical Physics, 1939. 7(12): p. 1103–1112.

20. Avrami, M., Kinetics of Phase Change. II Transformation-Time Relations for Random Distribution of Nuclei. The Journal of Chemical Physics, 1940. 8(2): p. 212–224.

21. Avrami, M., Kinetics of Phase Change. III Granulation, Phase Change, and Microstructure. The Journal of Chemical Physics, 1941. 9(2): p. 177–184.

22. Shirzad, K. and C. Viney, A critical review on applications of the Avrami equation beyond materials science. Journal of The Royal Society Interface, 2023. 20(203): p. 20230242.

23. Meijering, J., Interface area, edge length, and number of vertices in crystal aggregates with random nucleation. Philips Research Reports, 1953. 8: p. 270–290.

24. Todinov, M.T., A new approach to the kinetics of a phase transformation with constant radial growth rate. Acta Materialia, 1996. 44(12): p. 4697–4703.

25. Todinov, M.T., On some limitations of the Johnson–Mehl–Avrami–Kolmogorov equation. Acta Materialia, 2000. 48(17): p. 4217–4224.

26. Hunter, L., et al., Retrokinetics of crystallization. Scripta Materialia, 2025. 267: p. 116799.

27. Bertsekas, D. and J.N. Tsitsiklis, Introduction to Probability. 2nd ed. 2008, Belmont, MA: Athena Scientific.

28. Tomellini, M., On the grain size distribution function in KJMA compliant growth. Journal of Crystal Growth, 2022. 584: p. 126579.

29. Fanfoni, M. and M. Tomellini, Comment on: Retrokinetics of crystallization. Scripta Materialia, 2026. 280: p. 117329.

30. Cahn, J.W. and W.C. Hagel, Theory of the Pearlite Reaction, in Decomposition of austenite by diffusional processes, Z.D. Zackey and H.I. Aaronson, Editors. 1960, Interscience: New York. p. 131.

