## Supplementary Code for "Founder advantages in cell colony geometric organisation": staggered_seeding.m

```
clear
clc
close all

%% Multi colony growth on a 2D periodic square with uniform seeding
L_phys  = 24;                   % mm of system edge length
L        = 1.0;                 % normalised system edge length
v_phys   = 0.05;                % mm/h
v        = v_phys/L_phys;       % normalised growth speed

T_end    = 120;                 % end time of simulation
t        = linspace(0,T_end,100);   % seeding times
Nrun     = 1000;                  % Monte Carlo runs
makeGIF  = true;                % save GIFs
showN    = 1;                   % number of GIFs to save
fps      = 10;                  % GIF frame rate
nSide    = 500;                 % grid points per side (total ~ nSide^2)
rng(1); 

nCovTimes   = 100;                      % times at which to record coverage
tCov        = linspace(0,T_end,nCovTimes);
covFrac_all = zeros(Nrun, nCovTimes);   % store coverage for each run
K = numel(t);

% Sort times in increasing order and store inverse permutation
[ts,perm] = sort(t(:), 'ascend');
invperm   = zeros(K,1);
invperm(perm) = 1:K;

% 2D periodic grid on [0,L) x [0,L)
x = linspace(0, L, nSide+1);
x(end) = [];
[X, Y] = meshgrid(x, x);
dx     = L / nSide;
cellA  = dx^2;
nGrid  = numel(X);
W_all  = zeros(Nrun, K);

for run = 1:Nrun
    % Sample births uniformly across the system. Colonies born in
    % already transformed regions are kept as NaN and contribute zero area.
    xs = nan(K,1);
    ys = nan(K,1);
    for i = 1:K
        xi = L*rand;
        yi = L*rand;
    
        formed = ~isnan(xs) & (ts < ts(i));  % earlier accepted (non-phantom) seeds
        if any(formed)
            dxp = abs(xi - xs(formed));  dxp = min(dxp, L - dxp);
            dyp = abs(yi - ys(formed));  dyp = min(dyp, L - dyp);
            dist = hypot(dxp, dyp);
    
            r = v * (ts(i) - ts(formed)); % radii of earlier grains at time ts(i)
            if any(dist <= r)
                % Phantom: leave xs(i),ys(i) as NaN so it never competes
                continue
            end
        end
        % Not covered at birth (non-phantom)
        xs(i) = xi;
        ys(i) = yi;
    end
        xsEff   = xs.';   % 1 x K
        ysEff   = ys.';   % 1 x K
        tsEff   = ts.';   % 1 x K
    
        nanCols = isnan(xsEff); % seeds that never formed
        xsEff(nanCols) = 0;
        ysEff(nanCols) = 0;
    
        % Distances on periodic square
        xx = X(:); % nGrid x 1
        yy = Y(:); % nGrid x 1
        dxp = abs(xx - xsEff);
        dxp = min(dxp, L - dxp);
        dyp = abs(yy - ysEff);
        dyp = min(dyp, L - dyp);
    
        D = hypot(dxp, dyp); % nGrid x K
        T = D ./ v + tsEff; % arrival times field
        T(:, nanCols) = inf; % seeds that never formed
        [Tmin, owner] = min(T, [], 2);
        % Reshape for pixel plotting
        TminGrid  = reshape(Tmin,  nSide, nSide);
        ownerGrid = reshape(owner, nSide, nSide);
    
        % Overall covered area fraction over time (this run)
        % For each time in tCov, fraction = mean(points with arrival time <= t)
        covFrac_all(run,:) = mean(Tmin <= tCov, 1);
        covered_final = Tmin <= T_end;
        % Accumulate areas over covered cells only
        A_sorted = accumarray(owner(covered_final), cellA, [K 1], @sum, 0).'; % sorted-time order
        A_run    = A_sorted(invperm); % original t order
        W_all(run,:) = A_run;
    
        % Generate GIF
        if makeGIF && run <= showN
            fname = sprintf('2D_run%d.gif', run);
    
            if all(isnan(xs))
                tStart = 0;
            else
                tStart = min(ts(~isnan(xs)));
                if isempty(tStart) || tStart > T_end
                    tStart = 0;
                end
            end
            nFrames = 80;
            tvec    = linspace(tStart, T_end, nFrames);
            fig = figure('Color','w','Position',[100 100 500 500]);
            ax  = axes('Parent',fig);
            axis(ax,'equal');
            xlim(ax,[0 L]); ylim(ax,[0 L]);
            ax.XTick = []; ax.YTick = [];
            box(ax,'on'); hold(ax,'on');
            baseMap = turbo(256);
            idx = randperm(size(baseMap,1), K);
            cols = baseMap(idx, :);
            % Seed markers
            p = gobjects(K,1);
            for i = 1:K
                if isnan(xs(i)), continue; end
                p(i) = plot(ax, xs(i), ys(i), 'o', ...
                            'MarkerFaceColor', cols(i,:), 'MarkerEdgeColor','k', ...
                            'MarkerSize', 6, 'Visible','off');
            end
            % Pixel image placeholder (NaNs are transparent/uncovered)
            hImg = imagesc(ax, x, x, nan(nSide, nSide));
            set(ax, 'YDir', 'normal'); % make y increase upwards
            colormap(ax, cols); caxis(ax, [1 K]);
            set(hImg, 'AlphaData', false(nSide,nSide)); % start fully transparent
        for it = 1:numel(tvec)
            tt = tvec(it);
            % Reveal seeds
            for i = 1:K
                if ~isnan(xs(i)) && isgraphics(p(i)) && strcmp(get(p(i),'Visible'),'off') && tt >= ts(i)
                    set(p(i),'Visible','on');
                end
            end
            % Pixel update: show winning nucleus index where arrived by tt
            C = ownerGrid;
            C(TminGrid > tt) = NaN; % uncovered set as NaN
            set(hImg, 'CData', C, 'AlphaData', ~isnan(C));
            title(ax, sprintf('Run %d   t = %.3f (T_{end} = %.3f)', run, tt, T_end));
            drawnow;
            fr = getframe(fig);
            [A,map] = rgb2ind(frame2im(fr), 256);
            if it == 1
                imwrite(A, map, fname, 'gif', 'LoopCount', inf, 'DelayTime', 1/fps);
            else
                imwrite(A, map, fname, 'gif', 'WriteMode', 'append', 'DelayTime', 1/fps);
            end
        end
        close(fig);
        fprintf('Saved GIF: %s  |  areas (t = %.3g) = %s\n', fname, T_end, mat2str(A_run,4));
    end
end

figure('Position',[575	459	714	446])
tl = tiledlayout(2,3,'TileSpacing','compact','Padding','compact');
nexttile(1); hold on
binWidth = 0.005;
edges = 0:binWidth:max(W_all(:));
centers = edges(1:end-1) + binWidth/2;
% Compute histogram from pooled non-zero areas across all selected runs
nPlot = min(size(W_all,1),100);
W_pool   = W_all(1:nPlot,:);
nzValues = W_pool(W_pool ~= 0);
meanCounts = histcounts(nzValues,edges) / nPlot;
plot(centers,meanCounts,'r-','LineWidth',2)
% Analytical typical cell size distribution approximation
% https://doi.org/10.1016/j.jcrysgro.2022.126579
nAna = 5000;
Iphantom = numel(t) / (L^2 * (max(t) - min(t)));
omega    = (Iphantom*pi*v^2/3)^(1/3);
lambda   = sqrt(2*pi)*v/omega;
areaScale = lambda^2;
sEnd = omega * (T_end - min(t));
s    = linspace(0,sEnd,nAna).';
E0 = exp(-s.^3);
E1 = s .* E0;
C0 = -flipud(cumtrapz(flipud(s),flipud(E0)));
C1 = -flipud(cumtrapz(flipud(s),flipud(E1)));
ared = exp(s.^3) .* (C1 - s .* C0);
Aana = areaScale * ared;
dtau = [diff(s); 0] / omega;
wAna = Iphantom * L^2 .* exp(-s.^3) .* dtau;
bin = discretize(Aana,edges);
good = ~isnan(bin);
anaCounts = accumarray(bin(good),wAna(good),[numel(edges)-1 1],@sum,0).';
plot(centers,anaCounts,'k-','LineWidth',2)
xlabel('Colony size fraction at τ')
ylabel('Count')
legend('Sim. typical size','Analytical approximation','Location','northeast')
axis square
box on
hold off

nexttile(2); hold on
tcol = t(:);
simMean_EXi = mean(W_all,1,'omitnan').';
pred_EXi = EXieqn(tcol,T_end,v);
plot(tcol,simMean_EXi,'r-','LineWidth',2)
plot(tcol,pred_EXi,'k--','LineWidth',2)
xlim([0 T_end])
xlabel('Seeding time (hours)')
ylabel('Colony size fraction at τ')
legend('Sim. E[X_i]','Analytical E[X_i]','Location','northeast')
axis square
box on
hold off

nexttile(3); hold on
Wpos = W_all;
Wpos(Wpos <= 0) = NaN;
simMean_EXi_Xi = mean(Wpos,1,'omitnan').';
pred_EXi_Xi = EXi_Xieqn(tcol,T_end,v);
plot(tcol,simMean_EXi_Xi,'r-','LineWidth',2)
plot(tcol,pred_EXi_Xi,'k:','LineWidth',2)
xlim([0 T_end])
xlabel('Seeding time (hours)')
ylabel('Colony size fraction at τ')
legend('Sim. E[X_i|X_i > 0]','Analytical E[X_i|X_i > 0]','Location','northeast')
axis square
box on
hold off

nexttile(5); hold on
covFrac_mean = mean(covFrac_all,1,'omitnan');
total_EXi = zeros(size(tCov));
total_EXi_Xi = zeros(size(tCov));
for k = 1:numel(tCov)
    total_EXi(k) = sum(EXieqn(tcol,tCov(k),v));
    total_EXi_Xi(k) = sum(EXi_Xieqn(tcol,tCov(k),v));
end
dt = max(0,tCov(:).' - tcol);
A  = pi*(v*dt).^2;
avrami = 1 - exp(-sum(A,1));
plot(tCov,avrami,'k-.', ...
     tCov,covFrac_mean,'r-', ...
     tCov,total_EXi,'k--', ...
     tCov,total_EXi_Xi,'k:', ...
     'LineWidth',2)
legend('Avrami Eqn.', ...
       'Average Sim. Total', ...
       'Sum over E[X_i]', ...
       'Sum over E[X_i|X_i > 0]', ...
       'Location','eastoutside')
xlim([0 T_end])
xlabel('Time (hours)')
ylabel('Total fraction covered')
axis square
box on
hold off

save results.mat

function p = EXi_Xieqn(tvec, tau, c)
    tvec = tvec(:); n = numel(tvec);
    wpts = unique(tvec(tvec>0 & tvec<tau));
    s = integral(@(t) integrand_vec_2D(t,tvec,c), 0, tau, ...
        'ArrayValued',true,'RelTol',1e-12,'AbsTol',0,'Waypoints',wpts);
    E_ti = pi*(c*max(tvec - tvec.', 0)).^2;
    oneMinusVV = exp(sum(log(max(1 - E_ti, eps)),2));
    p = s(:) ./ max(oneMinusVV, eps);
end

function p = EXieqn(tvec, tau, c)
    tvec = tvec(:);
    wpts = unique(tvec(tvec>0 & tvec<tau));
    p = integral(@(t) integrand_vec_2D(t,tvec,c), 0, tau, ...
        'ArrayValued',true,'RelTol',1e-12,'AbsTol',0,'Waypoints',wpts);
    p = p(:);
end

function val = integrand_vec_2D(t, tvec, c)
    t = t(:);
    E = pi*(c*max(t - tvec.', 0)).^2;
    oneMinus = max(1 - E, 0);
    prodAll  = exp(sum(log(max(oneMinus, eps)),2));
    prodExcl = prodAll ./ max(oneMinus, eps);
    dE = 2*pi*c^2*(t - tvec.').*(t >= tvec.');
    val = (dE .* prodExcl).';
end
```
