## Supplementary Information for "Founder advantages in cell colony geometric organisation"

Table S1. Simulation results of the mean size fraction of selected colonies, indexed by their seeding time order $t_{n,i}$. Both the unconditional column mean and the column mean of the nonzero set are presented and compared with predictions from Eqn. 2 and Eqn. 3. Values are presented to 4 d.p.

| | $t_{n,1}$ | $t_{n,2}$ | $t_{n,3}$ | $t_{n,51}$ | $t_{n,52}$ | $t_{n,53}$ |
|---|---|---|---|---|---|---|
| Mean colony size fraction | 0.0512 | 0.0491 | 0.0474 | 0.0022 | 0.0020 | 0.0018 |
| Eqn. 2 | 0.0512 | 0.0492 | 0.0473 | 0.0022 | 0.0020 | 0.0018 |
| Mean of nonzero colony size fractions | 0.0512 | 0.0491 | 0.0474 | 0.0053 | 0.0050 | 0.0047 |
| Eqn. 3 | 0.0512 | 0.0492 | 0.0473 | 0.0052 | 0.0050 | 0.0047 |

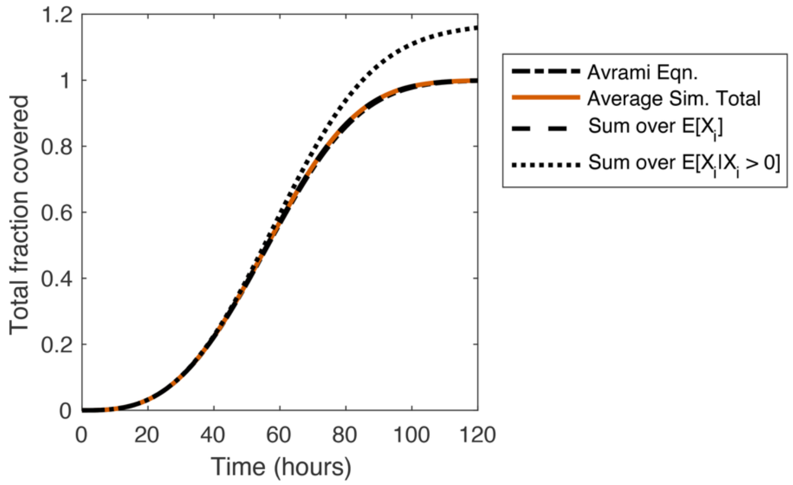


Figure S1. Total fraction covered over time: averaged sum over all colony sizes (red) compared to the prediction of the Avrami equation (Eqn. 1), the sum over the unconditional expectations of colony size (Eqn. 2), and the sum over the conditional expectations of colony size (Eqn. 3).
